# Bias and precision of parameter estimates from models using polygenic scores to estimate environmental and genetic parental influences

**DOI:** 10.1101/2020.08.11.246827

**Authors:** Yongkang Kim, Jared V. Balbona, Matthew C. Keller

## Abstract

In a companion paper (Balbona et al. (2020)), we introduced a series of causal models that use polygenic scores from transmitted and nontransmitted alleles, the offspring trait, and parental traits to estimate the variation due to the environmental influences the parental trait has on the offspring trait (vertical transmission) as well as additive genetic effects. These models also estimate and account for the gene-gene and gene-environment covariation that arises from assortative mating and vertical transmission respectively. In the current study, we simulated polygenic scores and phenotypes of parents and offspring under genetic and vertical transmission scenarios, assuming two types of assortative mating. We instantiated the models from our companion paper in the OpenMx software, and compared the true values of parameters to maximum likelihood estimates from models fitted on the simulated data to quantify the bias and precision of estimates. We show that parameter estimates from these models are unbiased when assumptions are met, but as expected, they are biased to the degree that assumptions are unmet. Standard errors of the estimated variances due to vertical transmission and to genetic effects decrease with increasing sample sizes and with increasing *r*^2^ values of the polygenic score. Even when the polygenic score explains a modest amount of trait variation (*r*^2^ = .05), standard errors of these standardized estimates were reasonable (< .05) for *n* = 16*K* trios, and smaller sample sizes (e.g., down to 4*K*) when the polygenic score is more predictive. These causal models offer a novel approach for understanding how parents influence their offspring, but their use requires polygenic scores on relevant traits that are modestly predictive (e.g., *r*^2^ > .025) as well as datasets with genomic and phenotypic information on parents and offspring. The utility of polygenic scores for elucidating parental influences should thus serve as additional motivation for large genomic biobanks to perform GWAS’s on traits that may be relevant to parenting and to oversample close relatives, particularly parents and offspring.

## Introduction

While behavioral genetics is sometimes viewed as being concerned with cataloging heritability (*h*^2^) and its determinants across traits, there has traditionally been great interest in understanding how family members directly impact each other environmentally. However, disentangling the genetic and environmental factors that cause familial resemblance has proven difficult, and is made all the more so when these factors are correlated. One likely reason for such a correlation is *vertical transmission* (VT), which occurs when a parental trait has a direct environmental influence on an offspring trait. VT leads to a covariance between the trait’s genetic and parental influences— a type of gene-environment covariance that has recently been termed *genetic nurture* by Kong et al. (2018).

Kong et al. (2018) showed that genetic nurture can be estimated from the covariance between the offspring’s phenotypic value and a polygenic score (PGS) calculated from the alleles not transmitted from parents to offspring. This covariance was denoted as *θ*_*NT*_ in Kong et al. (2018), which we also adopt here. Furthermore, the direct genetic effect of the PGS after removing the influence of genetic nurture can be estimated by subtracting *θ*_*NT*_ from the covariance between the transmitted PGS and the offspring phenotypic value (*θ*_*T*_). However, primary phenotypic assortative mating, which we denote in this paper simply as assortative mating (AM), occurs when mates choose each other based on phenotypic similarity, and complicates the modeling of *θ*_*NT*_ and *θ*_*T*_ considerably. A single generation of AM on a heritable trait leads to a positive ”trans” covariance between mates’ PGS’s (Robinson et al. (2017); Hugh-Jones et al. (2016)). Such a covariance is a competing explanation for non-zero observations of *θ*_*NT*_; specifically, part of *θ*_*NT*_ may be due to the AM-induced covariance between the transmitted PGS from the mother (which has a direct genetic influence on the offspring), the nontransmitted PGS from the father, and vice-versa.

Kong et al. recognized this confounding influence of AM on estimates of genetic nurture. In their study on educational attainment, they found evidence for AM in the parental generation but not before, and their approach to account for AM was therefore restricted to this scenario (disequilibrium AM). However, when AM occurs across multiple generations (equilibrium AM), recombination mixes causal variants of different parental origins on the same haplotype, leading to ”cis” covariance between causal variants within-haplotypes that is eventually (at equilibrium) equal to the trans covariance across mates. In this case, part of *θ*_*NT*_ may also be due to the AM-induced cis covariance between the transmitted PGS and the nontransmitted PGS from the same parent. Therefore, when there is evidence for equilibrium AM, this additional covariance must be accounted for to avoid bias.

In our companion paper (Balbona et al. (2020)), we introduce a series of causal models that use transmitted and nontransmitted PGS’s, along with the offspring phenotypic value, to estimate genetic nurture and direct genetic effects of the PGS under both equilibrium and disequilibrium AM scenarios. Importantly, these models also provide estimates of the full variance due to VT (*V*_*F*_). In particular, we showed that the full *V*_*F*_ can be estimated when there is no AM regardless of the predictive ability of the PGS 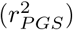. When there is AM, inclusion of parental phenotypic data or an assumption about 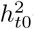 — the total *h*^2^ in the *base population* (before AM, or at ”time 0”)— allows estimates of the full *V*_*F*_, the full genetic nurture, and the full additive genetic variation. The causal modeling framework we used allows these models to be easily extended to account for different mechanisms of AM, to incorporate sibling and/or twin data, and to fit two traits bivariately in order to test cross-trait VT and AM.

The goal of this paper is to quantify the performance of the base models described in Balbona et al. (2020) when their assumptions are met, and to understand their sensitivity to assumptions when they are unmet. To do this, we simulated additive genetic effects, vertical transmission, and two types of AM (equilibrium and disequilibrium), and generated polygenic scores and phenotypes of parents and offspring. We instantiated the models in the OpenMx (Boker et al. (2011); Neale et al. (2016)) structural equation modeling software, and compared the true values of parameters to maximum likelihood estimates from the simulated data to quantify their bias and precision.

## Methods

### Causal models

In Balbona et al. (2020), we described three causal models that differ in their assumptions and data used, and we derive the expectations of parameters in each. Model 0 uses data on the offspring phenotype (*Y*_*o*_) and on PGS’s calculated from four sources: the transmitted paternal (*T*_*p*_), the nontransmitted paternal (*NT*_*p*_), the transmitted maternal (*T*_*m*_), and the nontransmitted maternal (*NT*_*m*_) haplotypes. Model 0 assumes no AM and no genetic effects other than those due to the PGS. Counter-intuitively, even when this last assumption is violated and the PGS explains little trait *h*^2^, we show mathematically that the estimate of the variance due to VT is unbiased. We verify this conclusion below. Note that VT is a process, not a score. In our path diagrams (Balbona et al. (2020)), we denote *F* as the ”familial” score caused by VT. Hence, the variation due to VT is denoted *V*_*F*_ with estimate 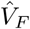

Model 1 uses the same data as Model 0, but incorporates the influence of AM on parameter expectations. As with Model 0, it assumes the PGS explains all genetic variation. Even though it models the influence of AM, Model 1 does not account for the influences of AM on the genetic effects not captured by the PGS. Thus, both Model 0 and Model 1 yield biased estimates when there is AM and the PGS explains less than the full *h*^2^. Because no PGS currently explains the full *h*^2^ for any trait, these two models should not be used when there is evidence of AM, and the utility of modeling AM in Model 1 is mostly a didactic example of how AM can be modeled.

In addition to the data used in the models above, Model 2 includes observed maternal (*Y*_*m*_) and paternal (*Y*_*p*_) phenotypic values in order to model the effects of both the PGS (with variance estimate 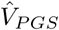) and the latent genetic score (LGS, with variance estimate 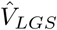). The LGS is the unobserved genetic score defined to be statistically orthogonal to the PGS in the base population. Furthermore, Model 2 provides estimates of the full genetic nurture effect 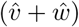, due to the covariance of parental effects with both the PGS 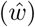 and the LGS 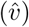. We also investigated the performance of a modification of Model 2 (Model 2-NP) that uses no parental phenotype information, and instead uses assumed values of the *h*^2^ in the base population, or at ”time 0” 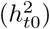. Such assumed values would presumably come from the literature, from models that provide decent estimates of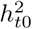 after accounting for AM and genetic nurture, such as extended twin family models (Keller et al. (2010)) or relatedness disequilibrium regression (RDR; Young et al. (2018); see Balbona et al. (2020) for a caveat about using estimates from RDR for this). For Models 1 and 2, we either assumed that AM has reached equilibrium (Models 1e and 2e) or that it is at disequilibrium (Models 1d and 2d), having occurred for only a single generation. Modeling other types of AM is discussed in our companion paper, but these are not examined here.

Details on model assumptions and parameter expectations for each model are in Balbona et al. (2020). Table 1 of Balbona et al. (2020) provides an overview of all parameter names, whereas Table 1 of this paper provides an overview of the principal differences between the models. We translated each of the seven models in Table 1 into OpenMx code in R. We used the NPSOL optimizer (Gill et al. (1986)) in OpenMx due to its ability to handle the many nonlinear constraints that were required to fit these models. The OpenMx scripts used here are available at https://github.com/yoki5348/VT_SEM.

**Table 1.** Description of causal models

| Model | AM | Use $Y_p$ and $Y_m$? | Assume $r_{PGS,t0}^2 = h_{t0}^2$? |
| --- | --- | --- | --- |
| 0 | no AM | No | Yes |
| 1e | equilibrium | No | Yes |
| 1d | disequilibrium | No | Yes |
| 2e | equilibrium | Yes | No |
| 2d | disequilibrium | Yes | No |
| 2e-NP | equilibrium | No | No |
| 2d-NP | disequilibrium | No | No |

### Data simulation

We simulated data using a modified version of the *GeneEvolve* software (Tahmasbi and Keller (2017)), which can simulate all the processes discussed here and create phenotypic data and genotypic data that has the same patterns of physical linkage disequilibrium observed in real SNP or sequence data. Our data did not require realistic patterns of linkage disequilibrium, and given that we needed to generate thousands of simulated datasets, we created a modified version of *GeneEvolve* in R in which all causal variants (CVs) were in linkage equilibrium in the base population. AM thereafter created realistic directional covariances between CVs.

For each simulation, we drew *m* = 100 binomially distributed CVs with minor allele frequencies (*p*) drawn from ~ *U* (.1, .5) and with effect sizes 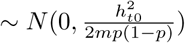. Half of the CVs contributed to the PGS and half to the LGS. We scaled the PGS so that it explained varying proportions of *V*_*Y*_ in the base population 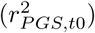, from 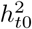 to .025, and we scaled the LGS to explain 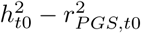 proportions of *V*_*Y*_. We summed the PGS and LGS to create a total genetic score and created an environmental score *ϵ* ∼ *N* (0, *V*_(*PGS*+*LGS*),*t*0)_ such that 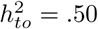 and standardized *Y* such that *V*_*Y,t*0_ = 1 exactly in the base population. The same scaling coefficients were then used across all generations, such that the variances of parameters could increase over their base population values as a consequence of VT or AM.

*GeneEvolve* chose mates such that the mate correlation, *r*_*mate*_, was either 0 or .25 across generations. Each mate couple had two offspring. The two haplotypes of mates recombined at random, leading to four haplotypes (transmitted vs. nontransmitted crossed by maternal vs. paternal origin) for each offspring. To simulate VT, the offspring familial environment was *F* = *f Y*_*p*_ + *f Y*_*m*_, where 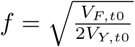 was constant across generations. It should be noted that *V*_*F*_ reaches equilibrium after a single generation, whereas AM takes ~ 5-10 generations before its consequences reach equilibrium. We therefore ran the modified *GeneEvolve* for a single generation to simulate disequilibrium AM, and for 20 generations to simulate equilibrium AM.

At the end of each simulation, we generated *n*_*fam*_ = 16*K* trio families such that no siblings existed in the final data, although more distant collateral relatives existed sporadically. Because our models only use within-family information to estimate parameters, we do not expect that non-independence across families due to distant relatives influenced our point estimates, but it may have led to slightly smaller standard errors (SE’s) of estimates than would occur if all families were unrelated. For the results presented in Figures 1, 2, and 3, 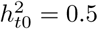, 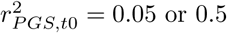, *V*_*F*,*t*0_ = 0.15, *r*_*mate*_ = 0 or 0.25, and AM was at equilibrium or disequilibrium. We also investigated other parameter values of 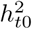, *r*_*mate*_, and *V*_*F,t*0_, but results from these simulations did not change conclusions and so for brevity, we do not present these results.

**Figure 1.**
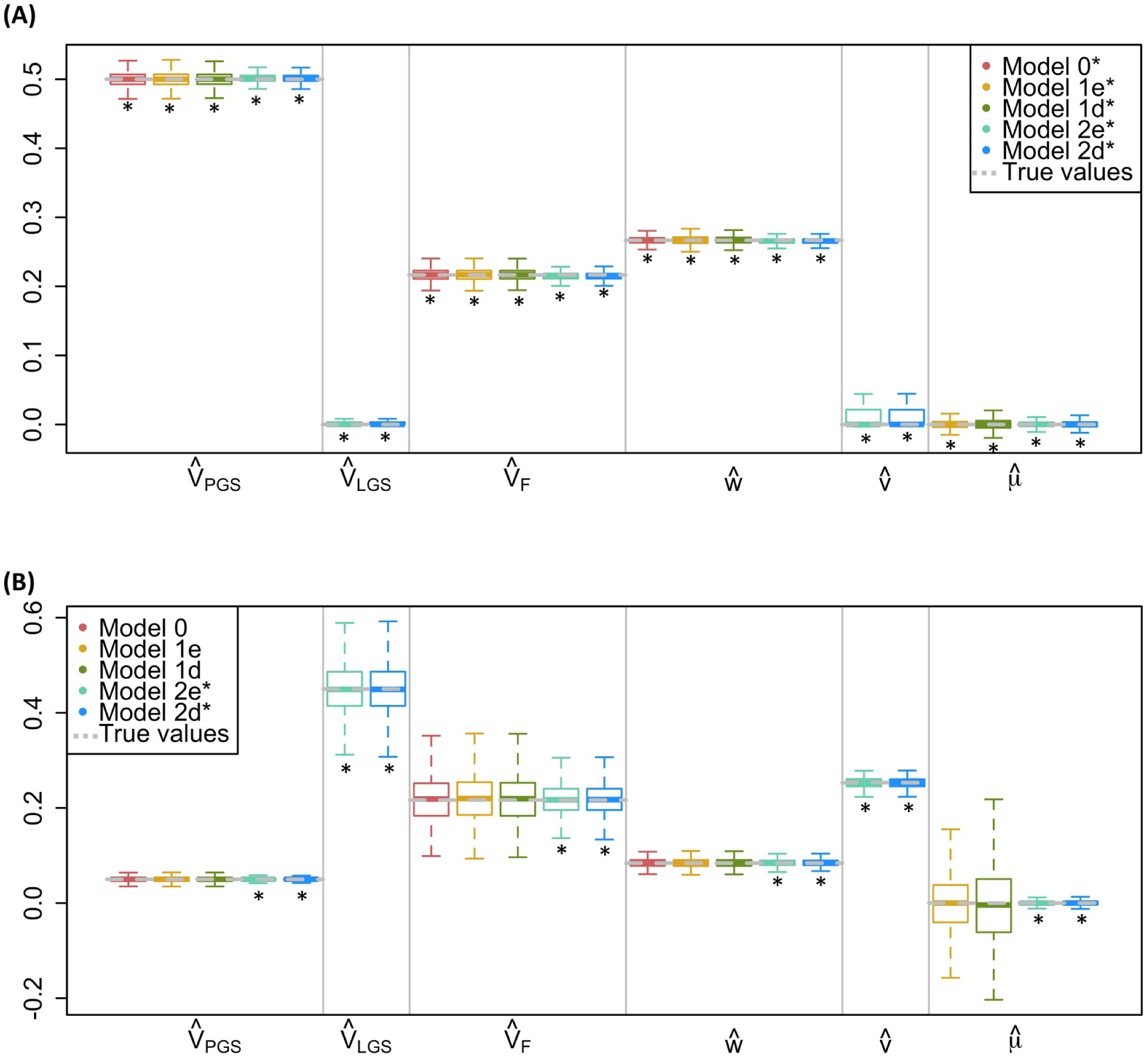
Comparison of estimates across models when there is VT but no AM. For each simulation, 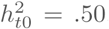, *r*_*mate*_ = 0, *V*_*F,t*0_ = .15, and *n*_*fam*_ = 16*K*. **(A)** 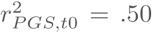. **(B)** 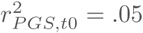. Boxplots show first quartile, median, and third quartile of estimates, with whiskers at the 2.5% and 97.5% quantiles. Equilibrium values of parameters are grey dashed lines. ∗ Models where assumptions about AM and 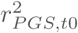 are met.

**Figure 2.**
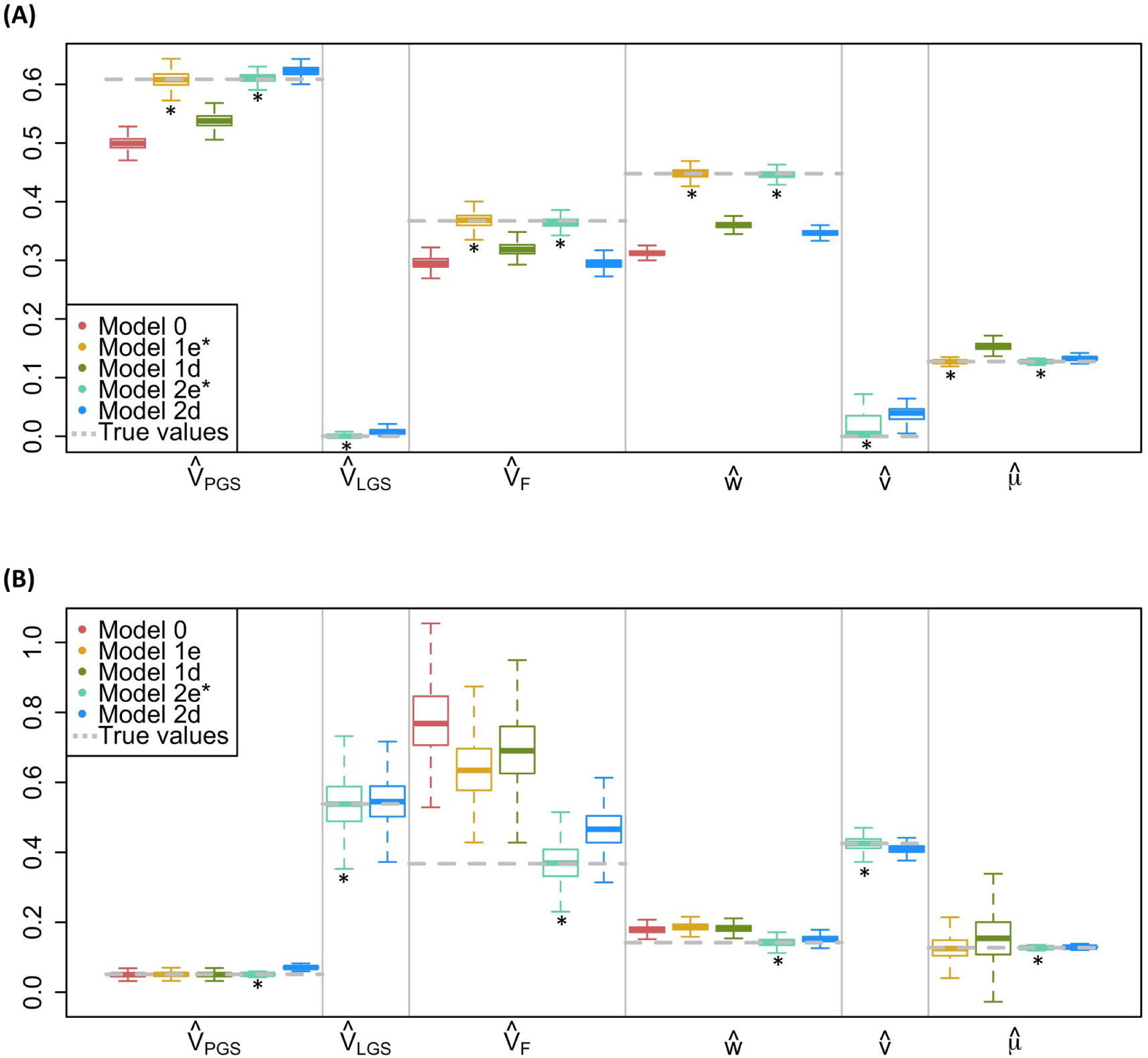
Comparison of estimates across models when there is VT and equilibrium AM. For each simulation, 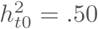, *r*_*mate*_ = .25, *V*_*F,t*0_ = .15, and *n*_*fam*_ = 16*K*. **(A)** 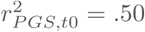. **(B)** 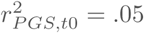. See Figure 1 note for additional details.

**Figure 3.**
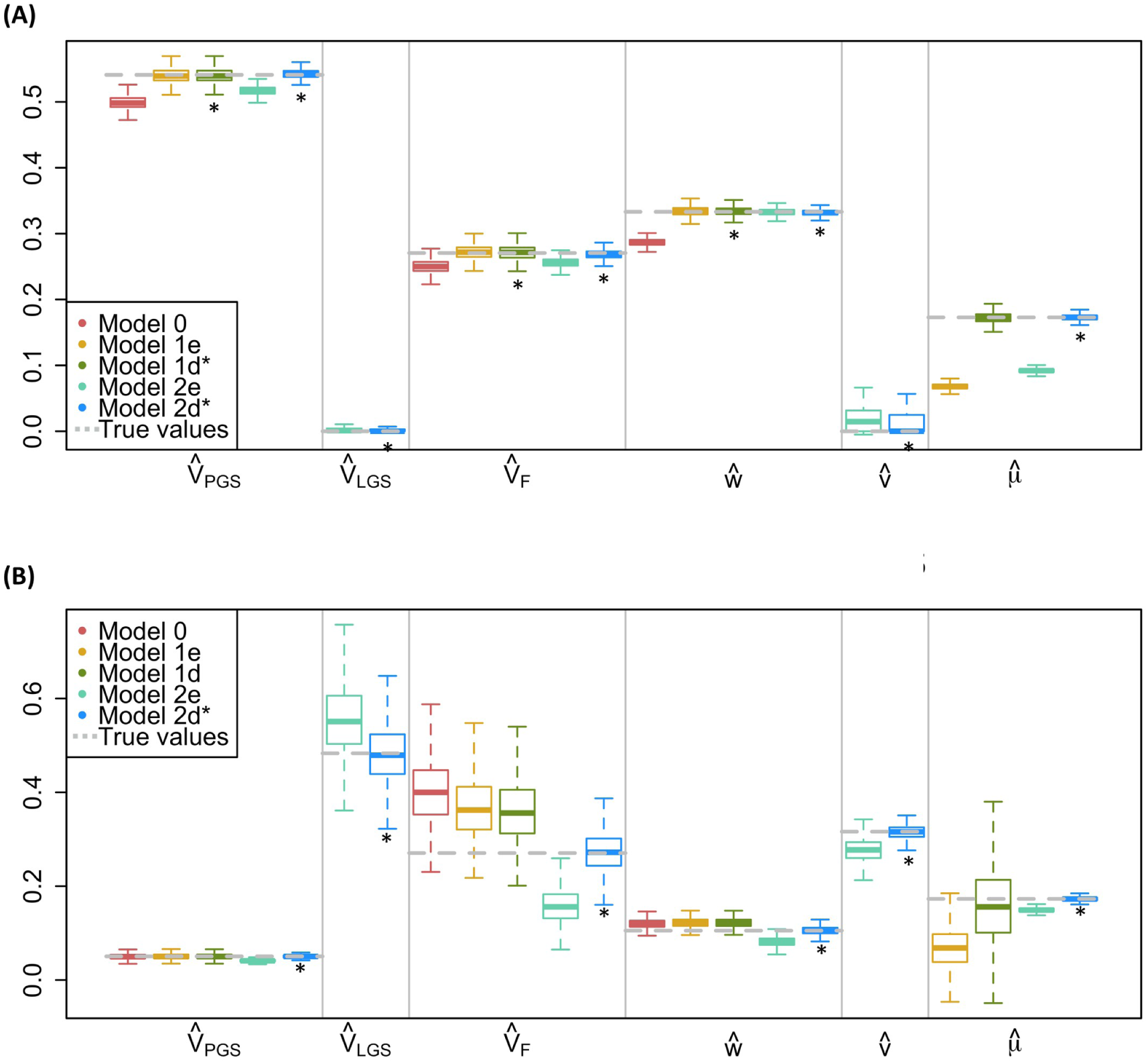
Comparison of estimates across models when there is VT and disequilibrium AM. For each simulation, 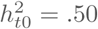, *r*_*mate*_ = .25, *V*_*F,t*0_ = .15, and *n*_*fam*_ = 16*K*. **(A)** 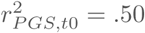. **(B)** 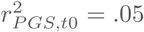. See Figure 1 note for additional details.

To estimate the standard errors for varying sample sizes, we generated datasets with the parameters 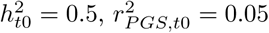, *r*_*mate*_ = 0.25, and *V*_*F,t*0_ = 0.15 and varied *n*_*fam*_ ∈ {1*K*, 2*K*, 4*K*, 8*K*, 16*K*, 32*K*, 64*K*}. To understand the influence of 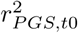, we used the above parameters but with fixed *n*_*fam*_ = 16*K* and varied 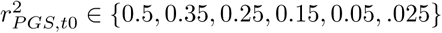. For all scenarios, we simulated data 1*K* times. So that the influence of AM and VT on the true parameter values are apparent, we chose not to standardize estimates in Figures 1–4, but we provide the equilibrium values of *V*_*Y*_ to aid in the interpretation of estimate values.

## Results

Figures 1, 2, and 3 show a comparison of estimates from five of the seven models we investigated. We show the results for the two models that used assumed values of 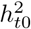 rather than parental phenotypes to derive estimates of latent genetic effects (Models 2e-NP and 2d-NP) in Supplement Figures 1-3. When the assumed values of 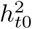 are correct, estimates from Models 2e-NP and 2d-NP are very similar to estimates from Models 2e and 2d, although their SE’s are slightly higher. Of course, estimates from Model 2e-NP and 2d-NP will be biased to the degree that assumed values of 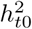 are incorrect. It should be noted that Model 2d-NP is most similar to the approach taken by Kong et al, although they did not attempt to estimate *V*_*F*_. Nevertheless, the results in Figures 1-3 from Model 2d should mimic most closely how their approach would perform, under the assumption that the value they assumed for 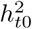 of educational attainment, taken from RDR (Young et al. (2018)), was correct.

Figures 1 shows the equilibrium true parameter values (grey dotted lines) and parameter estimates (boxplots) from the five models when there is no AM. In the absence of AM, all the models provide unbiased estimates of *V*_*PGS*_, *V*_*F*_, *w*, and *µ*, both when the PGS explains all the *h*^2^ (Figure 1A) or only 10% of it (Figure 1B). That Models 0 and 1 estimate the full *V*_*F*_ even when their assumption that 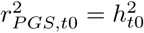 is violated confirms our conclusion in the companion paper, where we explain why this occurs. Thus, in the absence of AM, the full variation due to VT can be estimated simply from a weakly predictive PGS from both parents and values of the offspring trait. Model 2 provides unbiased estimates of *V*_*LGS*_ and *v*, and therefore provides full estimates of the additive genetic variation (*V*_*PGS*_ + *V*_*LGS*_) and the full genetic nurture effect (*w* + *v*). *V*_*LGS*_, *v*, and *µ* are not estimated in Model 0 and *V*_*LGS*_ and *v* are not estimated in Model 1, and so these estimates are not shown in the figures.

When there is equilibrium AM and the PGS explains all *h*^2^, the estimates from the models that assume equilibrium AM (1e and 2e) are unbiased (Figure 2A). Estimates from Model 2e are unbiased when there is equilibrium AM and the PGS is weakly predictive, but Model 1e’s estimates are sensitive to the assumption that 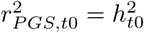 when AM exists (Figure 2B). This is because AM induces an unmodeled covariance between the PGS and the LGS (*i*) that inflates *θ*_*NT*_, which in turn upwardly biases estimates of *V*_*F*_ and *w* from Model 1e. However, Model 1e’s estimates of the direct effect of the PGS, which come from *θ*_*T*_ − *θ*_*NT*_, remain unbiased because *i* inflates *θ*_*NT*_ and *θ*_*T*_ to the same degree. Estimates from models that assume no AM (Model 0) or that model the wrong type of AM (Model 1d and 2d) do not properly account for the covariances between haplotypes that are induced by equilibrium AM, and therefore yield downwardly biased estimates when 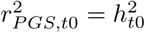 and upwardly biased estimates when 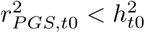. Nevertheless, the bias and spread of estimates from Model 2d are smaller than those of Model 1d because the observed mate covariance, as well as the covariance between one mate’s PGS and the other mate’s trait, are used in Model 2d, which decreases the bias in *µ* and therefore improves the estimation of other parameters.

When there has been a single generation of AM (disequilibrium AM), estimates from Model 2d are unbiased regardless of the predictive ability of the PGS, and estimates from Model 1d are unbiased as 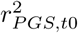 approaches 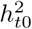 (Figure 3). When AM is at disequilibrium, estimates from models that assume equilibrium AM are typically biased. In particular, Model 2e’s estimates related to VT (*V*_*F*_, *w*, and *v*) are downwardly biased. This occurs because the covariances between haplotypic LGS’s and PGS’s (*g*, *h*, and *i*) implied by the causal model are larger than their actual values, which leads to expectations of *θ*_*NT*_ that are larger than those observed. To compensate, estimates of *V*_*F*_, *w*, and *v* are lowered while estimates of *V*_*LGS*_ are increased. Interestingly, when 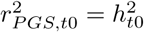, estimates of genetic nurture 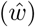 are unbiased when the model of AM is incorrect (Model 1e and 2e; Figure 3B). It is not obvious from the math why this this occurs, but may be because downwardly biasing 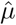 compensates for values of 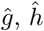, and 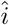 implied by the causal model that are too high.

The results for the disequilibrium AM scenario are similar to those from equilibrium AM in that they demonstrate the sensitivity of these models to assumptions about how AM has operated. Fortunately, as we describe in our companion paper, there is a good deal of information in the covariances between the four haplotypic PGS’s, the offspring trait, and potentially the two parental traits that allows assumptions regarding AM to be tested with high statistical power.

### Correlations and standard errors of parameter estimates

A causal model is considered ’under-identified’ when a set of two or more estimates use exactly the same information to estimate their values. All of the models we have reviewed above are identified. However, the information used to estimate different sets of parameters in these models is partially redundant, sometimes highly so, which decreases their precision. Figure 4 shows an example scatter plot between the estimates from the 1*K* simulated datasets from Model 2e (see Supplement Figure 4 for these results for Models 1e and 2e-NP). Much of the information to estimate *V*_*F*_ and *w* comes from *θ*_*NT*_, with both estimates increasing with higher values of *θ*_*NT*_, and so it is sensible that these two estimates are highly (*r* = .98) positively correlated. Furthermore, the assumption that the ratio of genetic nurture to direct genetic effects is the same for observed as for latent genetic effects 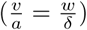 is required for Model 2 to be identified. Thus, 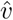 is an increasing function of 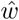 and therefore of *θ*_*NT*_; hence, 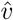 is positively correlated with 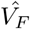 and 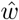. Much of the information to estimate *V*_*LGS*_ comes from the residual parent-offspring covariance, *cov*(*Y*_*o*_, *Y*_∗_), after removing the effects having to do with VT (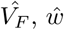, and 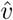), which explains the negative correlations between 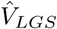 and these three estimates. Finally, 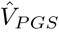 is not an observed variance (i.e., it is not simply synonymous with 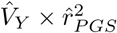). Rather, 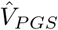 is the variance due to the *direct* effect of the PGS after removing its expected covariance with *F*, and so 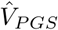 depends to some degree on the values of other estimates.

**Figure 4.**
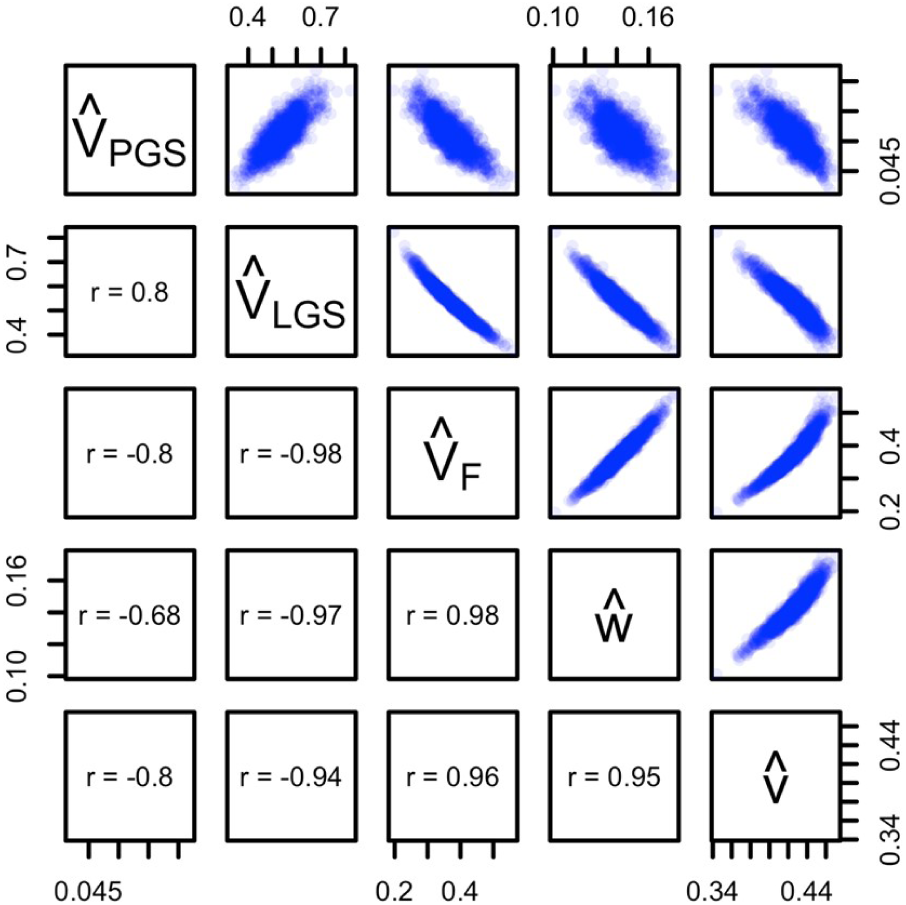
Scatter plots between Model 2e estimates. Estimates are from 1*K* simulations where 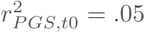, *r*_*mate*_ = 0.25, and AM is at equilibrium.

The high correlation values between parameter estimates suggests that their standard error (SE’s) will be high unless large sample sizes are employed. For the results displayed in Figure 5, we computed 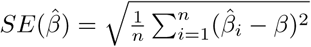, where 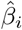 is the standardized estimate for simulated dataset *i*, and *β* is the true standardized value of the estimate. Note that unlike results in Figures 1–4, SE’s shown in Figure 5 are from standardized estimates for interpretability. Figure 5A shows the SE’s of estimates as a function of sample sizes (the number of trio families, *n*_*fam*_) from 1*K* to 64*K*. We compared the SE’s of models that provide unbiased estimates of *V*_*PGS*_, *V*_*F*_, and *w* when the PGS is weakly predictive and there is equilibrium AM (Models 2e and 2e-NP). Model 2e also estimates *V*_*LGS*_ whereas its value is assumed in Model 2e-NP, and so we show the SE’s of 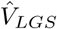 for Model 2e as well. As one would expect, SE’s of these estimates decrease as *n*_*fam*_ increases. Because their estimates correspond closely to observed statistics, 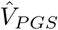 and 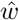 have much smaller SE’s than 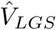 and 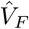. The SE’s of estimates from Model 2e are typically smaller than those from Model 2e-NP because Model 2e also uses information on parental phenotypes. To achieve a 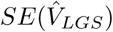 and 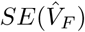 smaller than .05, at least *n*_*fam*_ = 8*K* trio families are required. These models can handle incomplete trios (only two of three family members sampled) so long as sufficient numbers of each type of relative pair exist in the sample. Of course, when there is such missingness, larger sample sizes are required. Finally, the SE’s of 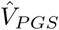 and 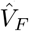 are slightly smaller for models that assume disequilibrium AM (Supplemental Figure 5), and so slightly smaller sample sizes are required for disequilibrium AM models to achieve equivalent statistical power.

**Figure 5.**
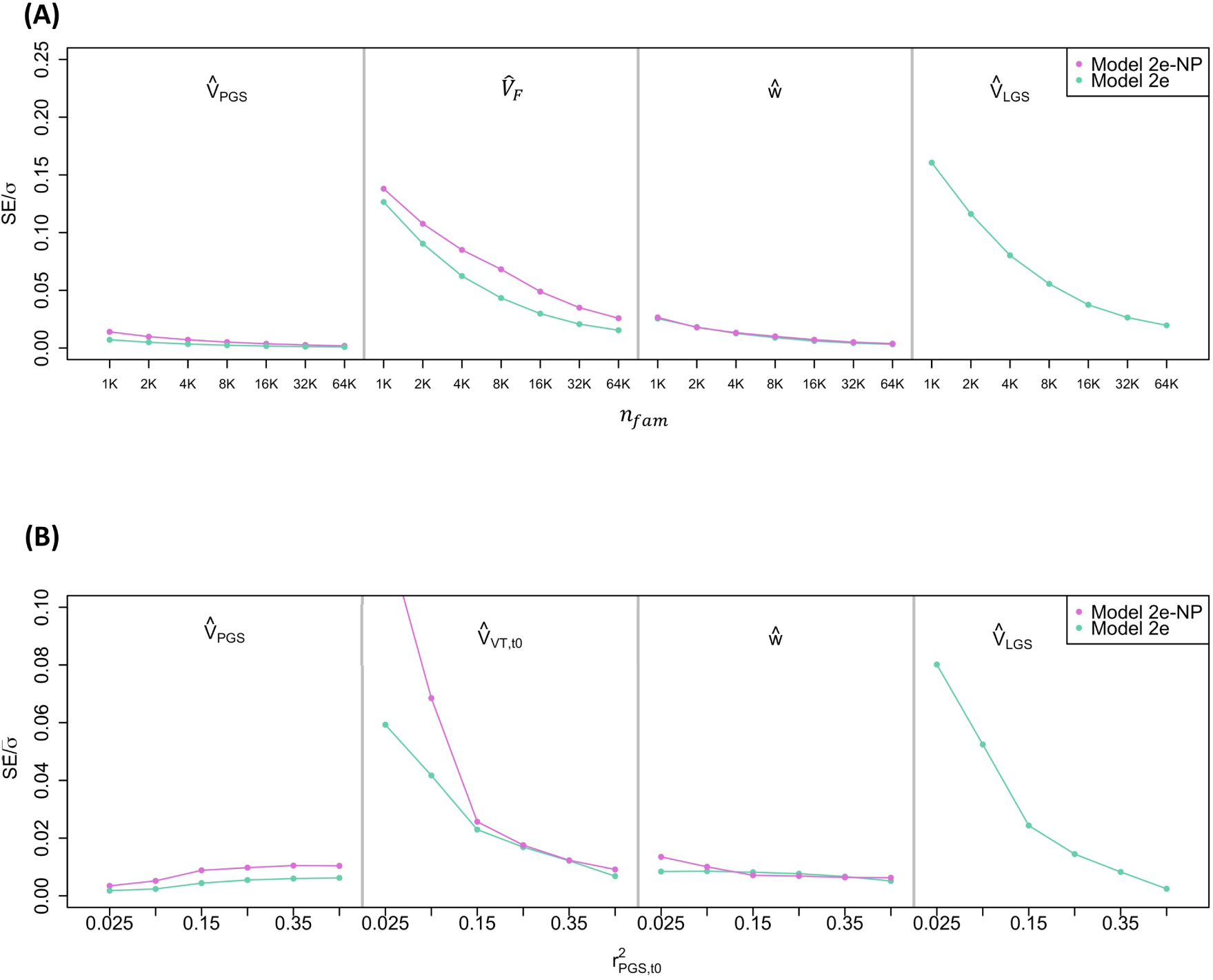
The standard errors (SE’s) of standardized estimates from Models 2e and 2e-NP. **(A) as a function of** *n*_*fam*_ **when** 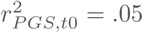 **and (B) as a function of** 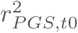 **when** *n*_*fam*_ = 16*K*. Estimates are from 1*K* simulations where *r*_*mate*_ = 0.25 and AM is at equilibrium.

Figure 5B shows the SE’s of estimates as a function of 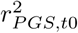 when *n*_*fam*_ is held constant at 16*K*. As 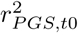 increases, the 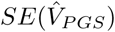 also increases slightly because the SE is proportionate to the mean value of the estimate. On the other hand, the 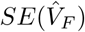 and 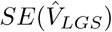 decrease as 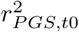 increases. This effect becomes more pronounced at low levels of 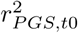, especially for Model 2e-NP. Information to estimate *V*_*F*_ comes primarily from 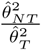 (Balbona et al. (2020)), and the variance of this ratio increases as 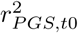 gets smaller. Because 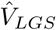 is strongly dependent on 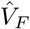 (Figure 4), the 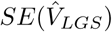 is similarly influenced by 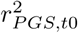. The relationships between 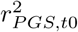 and the SE’s of estimates from models that assume disequilibrium AM are very similar to those shown (Supplemental Figure 5B).

### Computational performance

Table 2 shows the convergence time in seconds for each of the seven models for various sample sizes. We ran the models in OpenMx using the NPSOL optimizer with *feasibility tolerance*=1*e* − 7 and *Standard Errors* option set to *Yes*. For timing, we used a laptop with an i5 1.6GHz processor and 16GB RAM and ran each model and sample size combination a single time; hence there is some stochastic noise (e.g., Model 2e-NP took longer for *n*_*fam*_ = 4*K* than *n*_*fam*_ = 8*K*). As can be seen, despite their apparent complexity, the models run very fast. The slowest model (Model 2d) took only ~2.5 minutes with *n*_*fam*_ = 64*K*. Thus, computational capacity and time should not be limiting factors for using these models.

**Table 2.** Seconds until convergence across models.

| $n_{fam}$ | Model | | | | | | |
| --- | --- | --- | --- | --- | --- | --- | --- |
|  | 0 | 1e | 1d | 2e | 2d | 2e-NP | 2d-NP |
| 1K | 5.4 | 13.7 | 13.2 | 76.3 | 86.8 | 23.9 | 41.5 |
| 2K | 5.2 | 13.6 | 15.7 | 90.9 | 96.8 | 25.2 | 40.3 |
| 4K | 6.1 | 16.1 | 17.1 | 105.3 | 106.3 | 28.8 | 50.8 |
| 8K | 6.6 | 17.4 | 24.8 | 90.6 | 130.6 | 35.9 | 46.5 |
| 16K | 10.0 | 21.8 | 17.5 | 91.7 | 145.3 | 48.0 | 64.2 |
| 32K | 10.0 | 20.5 | 23.2 | 123.0 | 121.7 | 29.6 | 47.8 |
| 64K | 11.8 | 28.9 | 32.8 | 141.2 | 147.4 | 36.2 | 54.4 |

## Discussion

In this study, we quantified the performance of several causal models introduced in our companion paper that were inspired by Kong et al.’s approach for estimating genetic nurture. Using a Monte Carlo simulation approach to find true parameter values, generate trio datasets, and fit the simulated data in OpenMx, we confirmed that the estimates from these models are unbiased when assumptions are met. Indeed, when there is no AM, estimates from all models are unbiased even when the assumption regarding the predictive ability of the PGS is violated. However, when there is AM, estimates are sensitive to assumptions about the process leading to mate similarity; estimates are biased to the degree these assumptions are unmet. Fortunately, these assumptions do not have to be guessed at as the observed covariances involving the haplotypic PGS’s provide information that can be used to differentiate various processes of AM (Balbona et al. (2020)).

Model assumptions are never met perfectly in real data, and so the violation of an assumption does not mean that the estimates from the model are worthless, but it does mean that it is important to interpret the estimates with the proper nuance. It would be impossible to present results for estimates under all possible scenarios, but we cover some of the major ones in the figures above. In Table 3, we provide an overview of how violations of the principal assumptions influence parameter estimates, including some assumptions that were not covered in the present manuscript. We show the direction of the assumption violation that the biases refer to in the third column; the effect on the estimate(s) would be in the opposite direction for violations in the opposite direction. For example, *V*_*F*_ would be underestimated rather than overestimated if 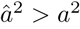. We omit the biases on 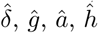, and 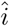 because their effects are already included in 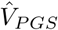 and 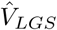. Of course, biases increase as violations become more severe, and so it is important to have some idea of how assumptions fare for any given data. When it is clear that assumptions are violated to a degree that makes estimates significantly biased, users should attempt to alter the model to accommodate new assumptions that better fit the data at hand.

**Table 3.**
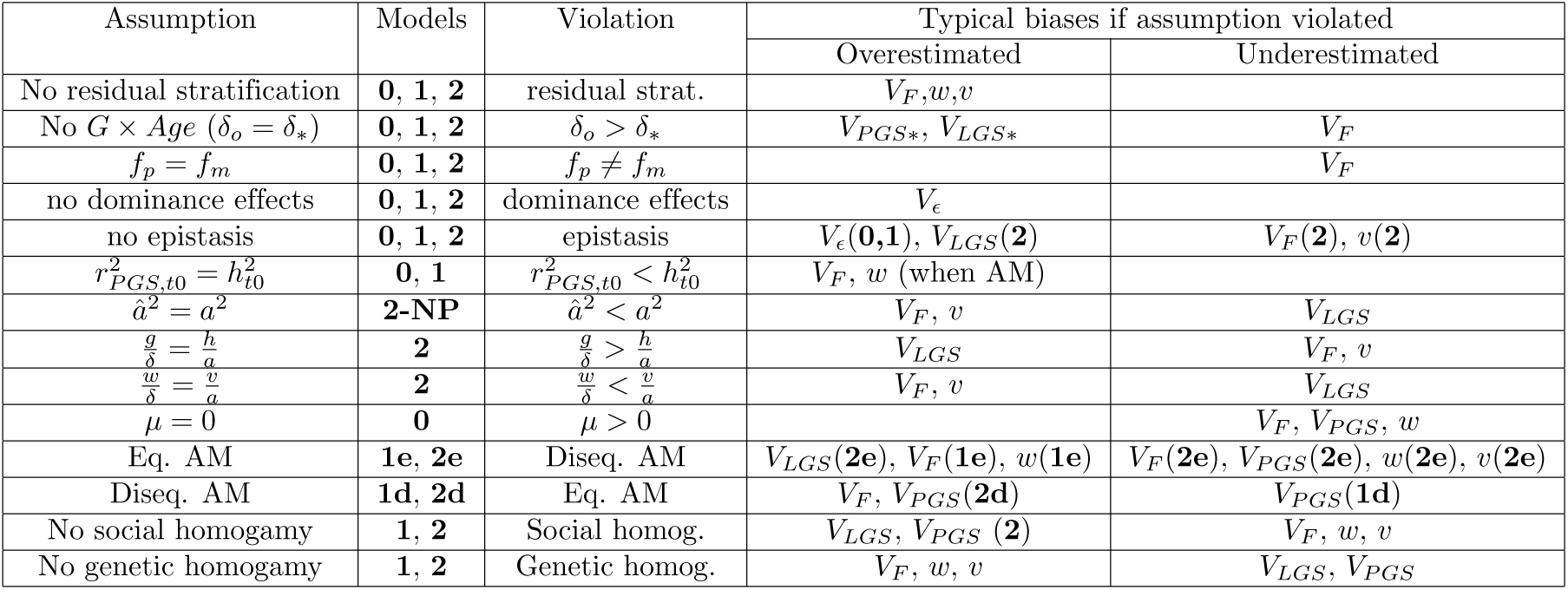
Effects of violating assumptions on parameter estimates.

Many of the influences of assumption violations covered in Table 3 are derivable from Figures 1–3 above. We briefly discuss two here that were not covered in our simulations. First, for some phenotypes, VT influences are likely to be stronger from one parent than the other (Kong et al. (2018)). When *f*_*p*_ ≠ *f*_*m*_, 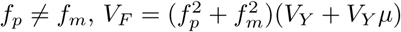. Using the current models, 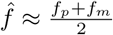. Thus, 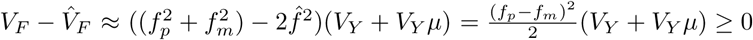. Therefore, *V*_*F*_ will be underestimated. Second, we discussed in our companion paper how different genetic effects in parents vs. offspring (*δ* ≠ *δ*_*o*_) can be modeled. When this is unmodeled, *δ*_*o*_ = *θ*_*T*_ − *θ*_*NT*_, as expected, and thus 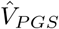 is unaffected (or minimally so when there is AM). However, *θ*_*NT*_ is a function of *δ*_∗_ rather than *δ*_*o*_, meaning that the observed *θ*_*NT*_ (=*cov*(*NT*_*m*_ + *NT*_*p*_, *Y*_*o*_)) will be smaller than that implied by the model, leading to underestimates of *V*_*F*_ and genetic nurture. Third, our models assume simple additive genetic effects with no dominance or epistasis. Because dominance does not inflate *cov*(*Y*_*o*_, *Y*_∗_), we expect that dominance influences would go only into the residual variance, 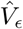. However, additive-by-additive (and higher order) epistasis would lead to unmodeled parent-offspring resemblance. For Model 2, this residual *cov*(*Y*_∗_, *Y*_*o*_) should upwardly bias 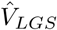 and have second-order effects on 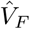 and 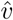.

The primary limitation to the current paper is that we made some of the same assumptions in our simulations as we did in our models, meaning that the influences of several factors on parameter estimates have yet to be investigated. We discuss a few of these in turn. First, both our simulations and our models assume no shared environmental influences, although we believe that there is sufficient information to estimate them. So long as there is no gene-environment covariance due to factors other than genetic nurture (e.g., stratification has been controlled for properly in the GWAS and in the structural equation models), *θ*_*NT*_ should only be influenced by genetic nurture and AM. Any residual differences between the mean phenotypic values of families after accounting for differences due to VT, genetic nurture, and genetics could be used to estimate shared environmental influences. Nevertheless, we have yet to build or test models that do this. Second, our models define the LGS as the genetic component that is statistically orthogonal to the PGS in the base population, and thus all covariance between the LGS and PGS (*i*) arises only from AM. Our simulations were based on this same assumption. We did not simulate physical linkage disequilibrium nor the process of building PGS’s based on estimated effects from GWAS. We do not anticipate any changes to our conclusions had we done this, but it is an issue that awaits confirmation. Finally, we have yet to simulate several scenarios (especially stratification, gene-by-age interactions, epistasis, social homogamy, and genetic homogamy) in Table 3, and so the reported influences on parameter estimates for these in Table 3 should be considered provisional until there is a more formal treatment.

The use of PGS’s to understand genetic nurture, as well as the direct genetic effect purged of their covariance with familial environmental effects, is an important advance made by Kong et al. We formalized this approach in a series of models introduced in our companion paper, and showed how this approach can also be used to estimate the total influence of parental traits on offspring traits. In the current paper, we have demonstrated that the models developed in our companion paper work as intended. These models are only the beginning, and they suggest many novel and exciting ways in which measured genetics data can be incorporated into family models to better understand the nature of nurture.

## Acknowledgments

We thank Rob Kirkpatrick for help with model fitting in OpenMx. This work was supported by grants R01MH100141 (to MCK) and T32MH016880 (to Dr. John Hewitt) and the Institute for Behavioral Genetics. This work utilized resources from the University of Colorado Boulder Research Computing Group, which is supported by the National Science Foundation (awards ACI-1532235 and ACI-1532236), the University of Colorado Boulder, and Colorado State University.

## References

Balbona, J., Kim, Y., and Keller, M. C. (2020). Estimation of parental effects using polygenic scores. (Under review).

Boker, S., Neale, M., Maes, H., Wilde, M., Spiegel, M., Brick, T., Spies, J., Estabrook, R., Kenny, S., Bates, T., Mehta, P., and Fox, J. (2011). Openmx: An open source extended structural equation modeling framework. Psychometrika, 76(2):306–317.

Gill, P. E., Murray, W., Saunders, M. A., Tomlin, J. A., and Wright, M. H. (1986). On projected newton barrier methods for linear programming and an equivalence to karmarkar’s projective method. Mathematical Programming, 36(2):183–209.

Hugh-Jones, D., Verweij, K. J., Pourcain, B. S., and Abdellaoui, A. (2016). Assortative mating on educational attainment leads to genetic spousal resemblance for polygenic scores. Intelligence, 59:103–108.

Keller, M. C., Medland, S. E., and Duncan, L. E. (2010). Are extended twin family designs worth the trouble? a comparison of the bias, precision, and accuracy of parameters estimated in four twin family models. Behavior genetics, 40:377–393.

Kong, A., Thorleifsson, G., Frigge, M. L., Vilhjalmsson, B. J., Young, A. I., Thorgeirsson, T. E., Benonisdottir, S., Oddsson, A., Halldorsson, B. V., Masson, G., Gudbjartsson, D. F., Helgason, A., Bjornsdottir, G., Thorsteinsdottir, U., and Stefansson, K. (2018). The nature of nurture: Effects of parental genotypes. Science, 359(6374):424–428.

Neale, M. C., Hunter, M. D., Pritikin, J. N., Zahery, M., Brick, T. R., Kirkpatrick, R. M., Estabrook, R., Bates, T. C., Maes, H. H., and Boker, S. M. (2016). Openmx 2.0: Extended structural equation and statistical modeling. Psychometrika, 81(2):535–549.

Robinson, M. R., Kleinman, A., Graff, M., Vinkhuyzen, A. A., Couper, D., Miller, M. B., Peyrot, W. J., Abdellaoui, A., Zietsch, B. P., Nolte, I. M., et al. (2017). Genetic evidence of assortative mating in humans. Nature Human Behaviour, 1(1):1–13.

Tahmasbi, R. and Keller, M. C. (2017). Geneevolve: a fast and memory efficient forward-time simulator of realistic whole-genome sequence and snp data. Bioinformatics (Oxford, England), 33:294–296.

Young, A. I., Frigge, M. L., Gudbjartsson, D. F., Thorleifsson, G., Bjornsdottir, G., Sulem, P., Masson, G., Thorsteinsdottir, U., Stefansson, K., and Kong, A. (2018). Relatedness disequilibrium regression estimates heritability without environmental bias. Nature Genetics, 50(9):1304–1310.

