## Supplement Figures and Tables for "Bias and precision of parameter estimates from models using polygenic scores to estimate environmental and genetic parental influences"

### (Supplementary Notes) Vertical transmission estimation via structural equation modeling (VT-SEM)

**1 Institute for Behavioral Genetics, University of Colorado at Boulder**

**2 Department of Psychology & Neuroscience, University of Colorado at Boulder**

*** ***

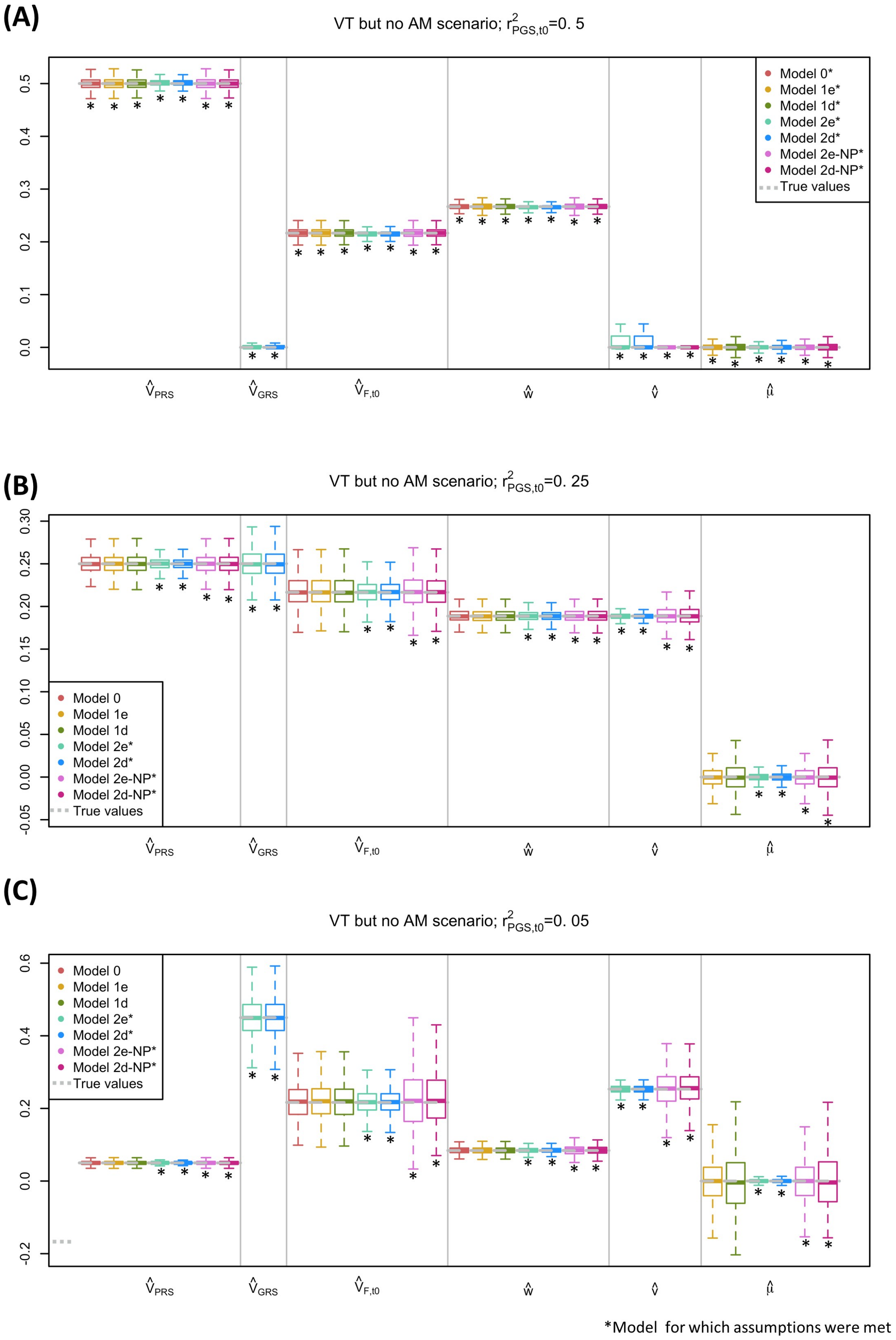


**Suppl Figure 1. Comparison of estimates across models when there is VT but no AM.** For each simulation, $h^2_{t0} = .50$, $r_{mate} = 0$, $V_{F,t0} = .15$, and $n_{fam} = 16K$. **(A)** $r^2_{PGS,t0} = .50$. **(B)** $r^2_{PGS,t0} = .25$. **(C)** $r^2_{PGS,t0} = .05$. Boxplots show first quartile, median, and third quartile of estimates, with whiskers at the 2.5% and 97.5% quantiles. Equilibrium values of parameters are grey dashed lines. $*$ Models where assumptions about AM and $r^2_{PGS,t0}$ are met.

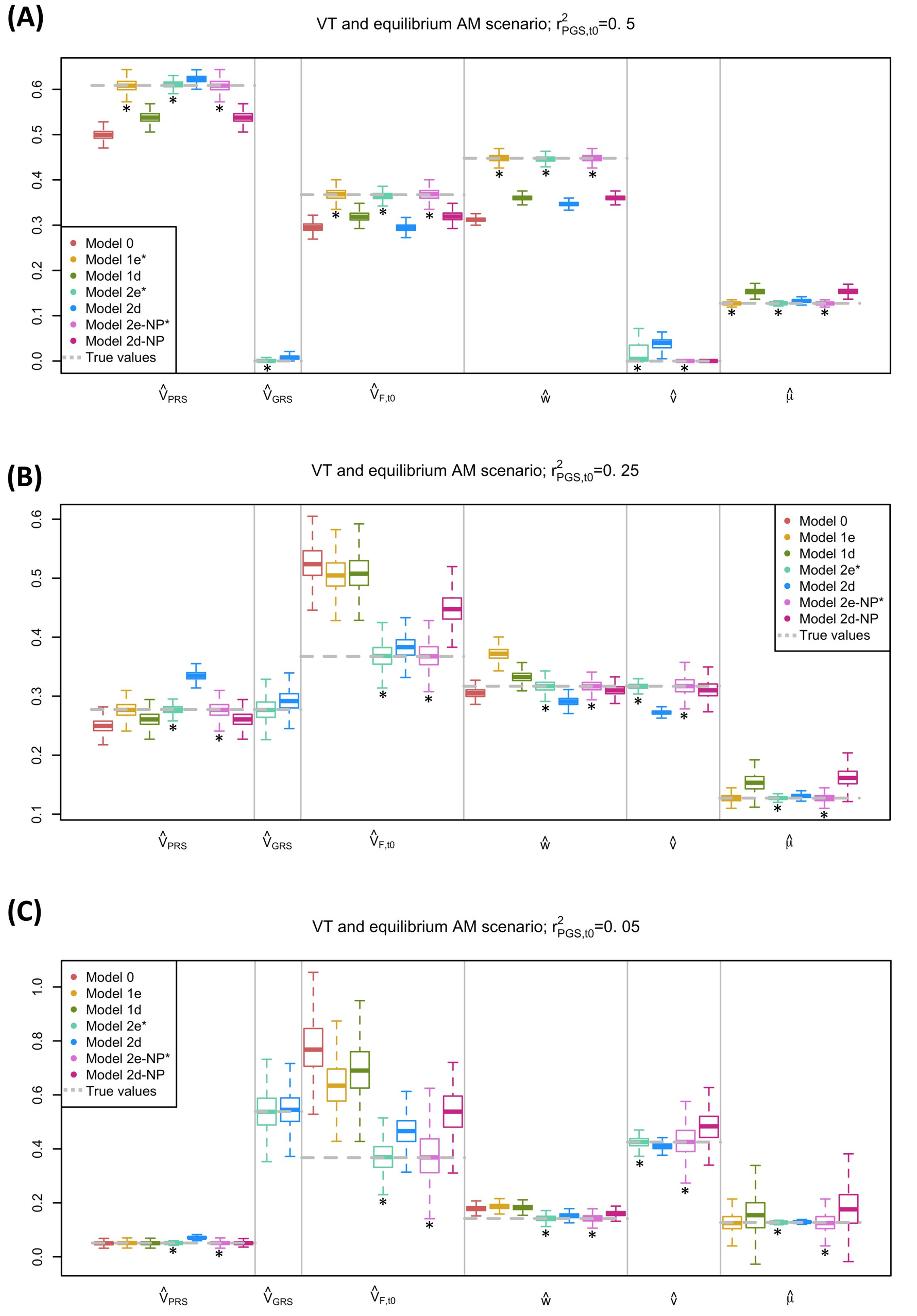


*Model for which assumptions were met

**Suppl Figure 2. Comparison of estimates across models when there is VT and equilibrium AM**. For each simulation, $h^2_{t0} = .50$, $r_{mate} = .25$, $V_{F,t0} = .15$, and $n_{fam} = 16K$. **(A)** $r^2_{PGS,t0} = .50$. **(B)** $r^2_{PGS,t0} = .25$. **(C)** $r^2_{PGS,t0} = .05$. See **Figure ??** note for additional details.

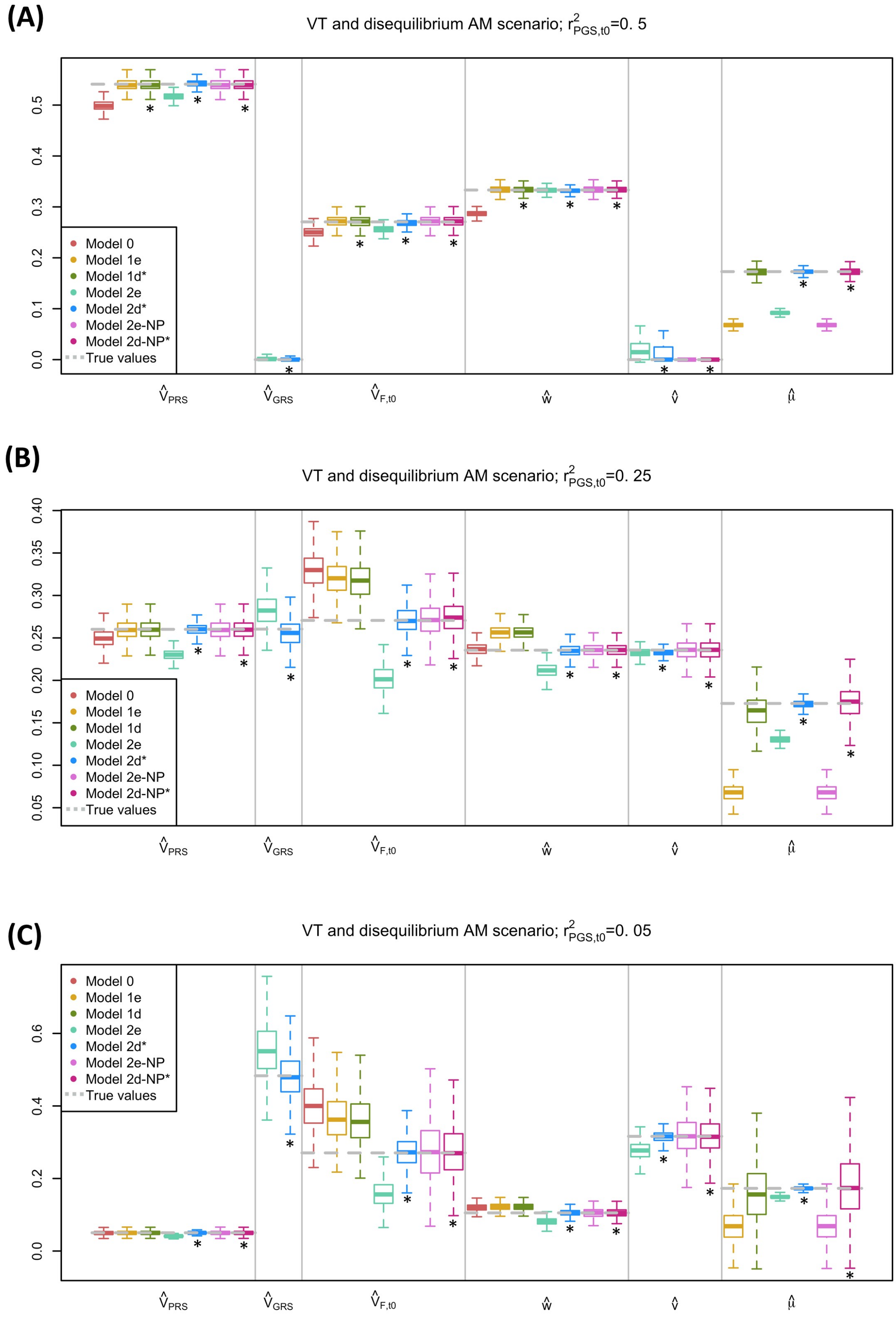


*Model for which assumptions were met

**Suppl Figure 3. Comparison of estimates across models when there is VT and disequilibrium AM**. For each simulation, $h^2_{t0} = .50$, $r_{mate} = .25$, $V_{F,t0} = .15$, and $n_{fam} = 16K$. **(A)** $r^2_{PGS,t0} = .50$. **(B)** $r^2_{PGS,t0} = .25$. **(C)** $r^2_{PGS,t0} = .05$. See **Figure ??** note for additional details.

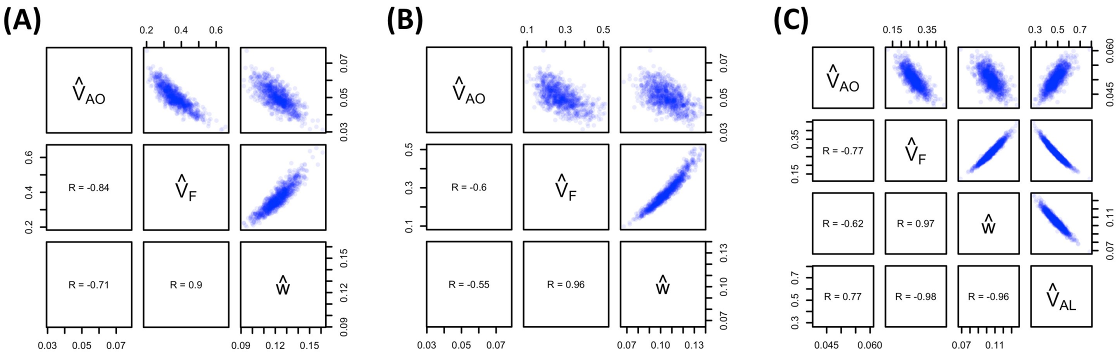


**Suppl Figure 4. Scatter plots between estimates**. Estimates are from $1K$ simulations where $r^2_{PGS,t0} = .05$, $r_{mate} = 0.25$, and AM is at disequilibrium. (A) shows scatter plots between estimates from model 1d. (B) shows scatter plots between estimates from model 2d-NP. (C) shows scatter plots between estimates from model 2d.

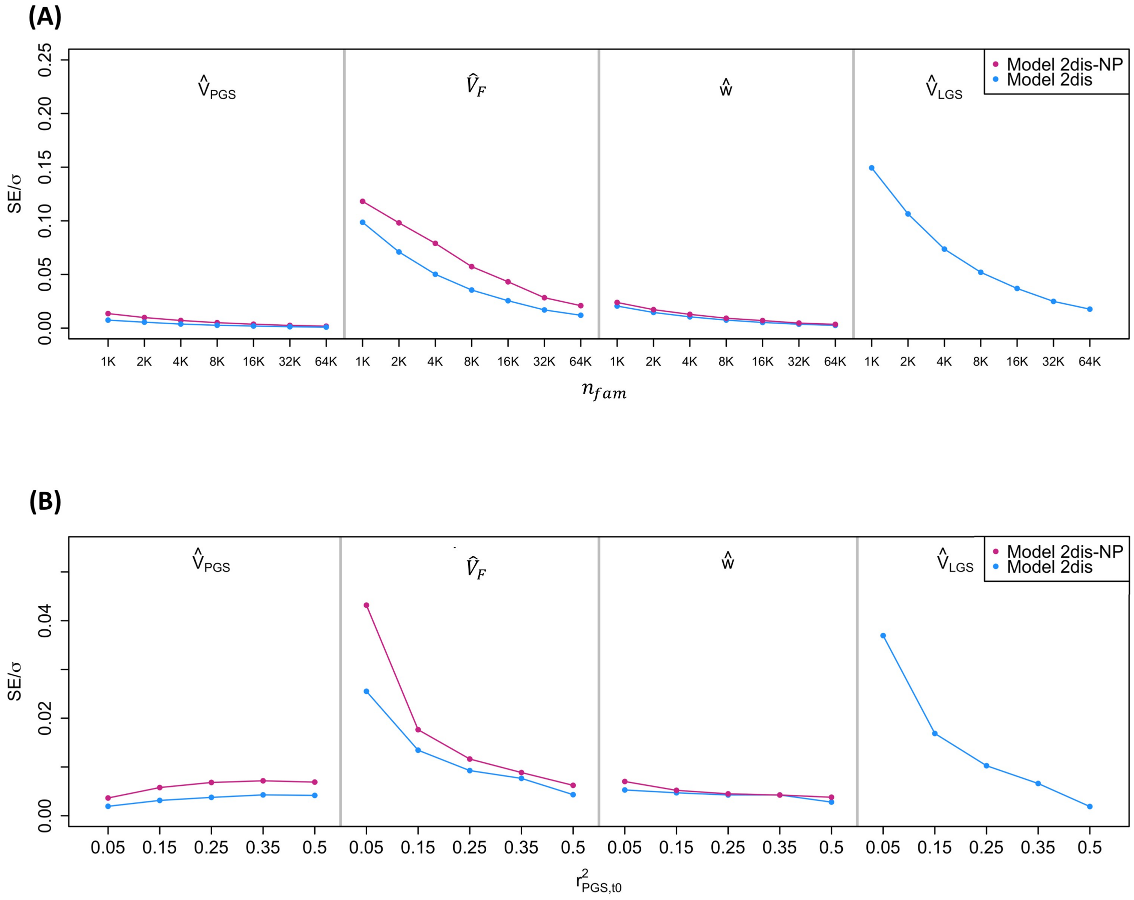


**Suppl Figure 5. The standard errors (SE's) of standardized estimates from Models 2e and 2e-NP (A) as a function of $n_{fam}$ when $r^2_{PGS,t0} = .05$ and (B) as a function of $r^2_{PGS,t0}$ when $n_{fam} = 16K$.** Estimates are from $1K$ simulations where $r_{mate} = 0.25$ and AM is at disequilibrium.
